# Skip-Zeros Variational Inference in the Million-Cell Era of Single-Cell Transcriptomics

**DOI:** 10.64898/2026.02.20.706458

**Authors:** Ko Abe, Shintaro Yuki, Teppei Shimamura

## Abstract

Combinatorial indexing-based single-cell RNA sequencing methods such as sci-RNA-seq and sci-RNA-seq3 now enable the profiling of millions of cells, producing expression matrices that are both extremely sparse and high-dimensional. Conventional nonnegative matrix factorization (NMF) provides an interpretable framework for uncovering latent biological structures but is computationally prohibitive at this scale, as it requires explicit access to the vast number of zero entries. We introduce UNISON (Unified Sparse-Optimized Nonnegative factorization), a scalable framework for matrix and tensor factorization based on skip-zeros variational inference. By reformulating stochastic variational Bayes updates in terms of sufficient statistics, UNISON performs inference using only nonzero elements, while implicitly accounting for zeros through geometric sampling. This strategy enables efficient parameter estimation without matrix expansion and naturally accommodates multiple experimental contexts. Simulation studies show that UNISON is robust to diverse learning-rate schedules and mini-batch sizes, providing practical guidelines for optimization. Application to the Mouse Organogenesis Cell Atlas demonstrates scalability to over one million cells, yielding latent factors that capture developmental trajectories and lineage-specific signatures with improved interpretability compared to existing methods. Cross-species analysis of aqueous humor outflow pathways across five vertebrate species further highlights UNISON’s ability to disentangle conserved from species-specific transcriptional programs and to recover biologically meaningful gene-gene and gene-phenotype relationships relevant to glaucoma. By efficiently exploiting sparsity while preserving interpretability, UNISON establishes a principled and practical solution for integrative, large-scale single-cell transcriptomics.

**Significance Statement:** Single-cell technologies now generate datasets spanning millions of cells, creating massive, sparse matrices that are computationally difficult to analyze. Conventional methods often sacrifice statistical rigor for speed, either by discarding data or employing inappropriate models. We present UNISON, a framework that leverages a mathematical trick to perform exact inference using only nonzero elements. By combining this efficient computation with a probability model tailored for count data, UNISON scales to millions of cells while preserving information on rare cell types. This approach resolves the trade-off between scalability and accuracy, enabling precise integrative analyses of development and evolution across species without requiring massive computational resources.

---

Recent technological advancements, such as sci-RNA-seq and sci-RNA-seq3 that leverage combinatorial indexing, have enabled the generation of massive single-cell libraries in a single experiment (e.g. 1). This has led to the acquisition of gene expression datasets comprising millions of cells, which are typically represented as high-dimensional and extremely sparse matrices.

Matrix factorization is a widely used statistical method for projecting high-dimensional matrices into a lower-dimensional space to extract latent patterns. Among these methods, Non-negative Matrix Factorization (NMF; 2, 3) is particularly advantageous due to its non-negativity constraints, which enhance the interpretability of the results. Consequently, NMF has become a crucial tool for analyzing high-dimensional biological data.

However, conventional NMF requires explicit handling of zero elements even in sparse matrices, and its application to datasets comprising millions of cells has often been hindered by the enormous computational cost and memory consumption. While methods such as UMAP (4) and t-SNE (5) are widely employed for dimensionality reduction, matrix factorization remains indispensable for several reasons. UMAP and t-SNE produce non-linear, stochastic embeddings with reproducibility issues and high parameter sensitivity (6), making them unsuitable for rigorous interpretation or extraction of reusable latent factors. In contrast, matrix factorization naturally connects to statistical models, enabling direct use of latent factors in downstream analyses such as gene network inference and cell population comparisons. Furthermore, being grounded in linear algebra, it provides computational efficiency and scalability for large-scale data analysis. Meanwhile, online NMF methods such as Liger (7) have been developed for large-scale data, achieving speedup through subsampling. However, using only a portion of the data may compromise statistical fidelity, and many of these methods optimize a Gaussian likelihood (i.e. mean squared error), which is inappropriate for discrete count data. In light of this background, effective analysis of noisy, high-dimensional single-cell data requires significant improvement in computational efficiency while maintaining statistical rigor, adoption of probability models appropriate for count data, and integrative utilization of prior biological knowledge.

In this study, we present UNISON (Unified Sparse-Optimized Nonnegative factorization), a methodological frame-work tailored to the million-cell era of single-cell transcriptomics. The core innovation of UNISON is “skip-zeros variational inference,” which performs parameter estimation using only nonzero elements, while implicitly accounting for the contribution of zero elements through sampling based on the geometric distribution. This enables direct manipulation of sparse matrices and allows analysis of datasets exceeding one million cells without expanding zero elements. Our primary contributions are as follows:

i. From an algorithmic perspective, we revisit stochastic gradient ascent (SGA) and its Bayesian counterpart, stochastic variational Bayes (SVB), which are widely used for large-scale parameter estimation. Naive implementations of these methods on sparse data are inefficient, as they require explicit expansion of zero elements. We resolve this inefficiency by reformulating SVB updates in terms of sufficient statistics, which enables parameter inference without accessing zero entries.
ii. Furthermore, UNISON framework can be applied to not only NMF but also unified nonnegative matrix factorization (UNMF; 8), which separates data format from experimental context and enables incorporation of background information such as species, batch, and experimental conditions through a design matrix. We derive an estimation procedure for UNMF that leverages the same skip-zeros principle, allowing complex analyses—such as integrating data across multiple species to disentangle conserved programs from species-specific variation—to be executed without compromising computational efficiency.

The remainder of this article is organized as follows. Results section presents overview of our methods, simulation studies, and applications to real single-cell datasets, demonstrating decomposition of more than one million cells as well as cross-species analysis integrating sequence-based background information.

## Results

### Practical implementation of the skip-zeros SVB framework

The methodological developments converge into *Unified Sparse-Optimized Nonnegative factorization* (UNISON). This frame-work encompasses both the standard NMF and its generalization to UNMF, and is designed to operate natively on sparse matrices expressed in coordinate (COO) format. In practice, the algorithm proceeds by repeatedly sampling mini-batches of nonzero entries from the sparse representation, computing sufficient statistics for these observations, and updating the variational parameters of the factor matrices. Because the sufficient statistic *T* (*y*) vanishes when *y* = 0, all updates can be expressed entirely in terms of nonzero elements. The contribution of zero entries, which dominate single-cell data, is incorporated implicitly through the use of a geometric distribution. This procedure ensures that the statistical effect of zeros is accounted for, without requiring their explicit enumeration. Importantly, because the algorithm never reconstructs the dense matrix, memory usage remains manageable even for datasets containing millions of cells, and large mini-batch sizes can be employed without incurring prohibitive computational cost. The key innovation of the framework is that it unifies generic stochastic variational Bayesian (SVB; 9, 10) method for sparse matrix factorization, Poisson NMF with skip-zeros, and UNMF with skip-zeros into a single coherent methodology. By operating directly on COO-formatted data—compatible with widely used preprocessing pipelines such as Cell Ranger—the framework integrates seamlessly into existing single-cell analysis workflows. Overall, UNISON, built upon skip-zeros variational inference, provides a practical and scalable solution for analyzing high-dimensional, noisy, and extremely sparse single-cell datasets, thereby addressing the challenges of the million-cell data era. It combines the interpretability of nonnegative factorization with the scalability of variational inference, thereby enabling analyses that were previously computationally infeasible, such as decomposition of over one million cells or integration of cross-species datasets within a unified latent space. See Materials & Methods for technical details.

### Simulation study

We first conducted simulation experiments to evaluate the statistical properties and computational behavior of the proposed skip-zeros SVB framework. The primary objectives were to examine how estimation accuracy and stability depend on mini-batch size and learning-rate parameters, and to highlight the computational advantages of omitting zero entries. Synthetic data were generated from a probabilistic generative process consistent with Poisson factorization. Specifically,

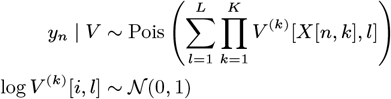

with the latent dimension fixed at *L* = 5. Here, the (*i, j*) element of any matrix *X* be written as *X*[*i, j*], and let the *j* th column vector of a matrix be denoted as *X*[:, *j*]. Two scenarios were considered: matrix factorization (*K* = 2) and tensor factorization (*K* = 3). Because decomposition is not unique up to scaling and permutation, components *V* [:, 1], …, *V* [:, *L*] were ordered by increasing variance, and recovery was evaluated using correlation coefficients that are invariant to scale.

For matrix factorization, the number of rows was fixed at 100, and the number of columns was set to 100, 500, or 1000. For tensor factorization, we considered *D*_1_ = 50, *D*_2_ = 500 or 1000, and *D*_3_ = 2. To assess robustness, we varied the following hyperparameters: mini-batch sizes of 500, 1000, and 2000; delay parameter *τ* ∈ {1.5, 5, 15} in the learningrate schedule; and forgetting rates *κ* ∈ {0.7, 0.8, 0.9}. The results, summarized in Figure 1, show that the skip-zeros SVB algorithm consistently recovered the latent structure across settings. Estimation stability and convergence speed, however, depended on dataset size and learning-rate scheduling. For smaller matrices, smaller delay values *τ* produced more stable estimates by enabling faster adaptation. For larger matrices, larger *τ* values were preferable, as slower decay of the learning rate ensured sufficient updates before convergence. Increasing mini-batch size generally improved stability, reflecting reduced variance in stochastic updates. Importantly, because zero entries were omitted, computational cost remained practical even for large mini-batch sizes, which would otherwise be infeasible in conventional implementations. These simulations provide clear guidance for practical use. For moderate-sized datasets, a small delay *τ* combined with a moderate forgetting rate (*κ* ≈ 0.8) is effective. For large-scale datasets with millions of cells, a larger delay (*τ* ≥ 15) is recommended to maintain stability. The robustness of the algorithm across a broad range of conditions confirms that zeros can be safely omitted without biasing inference.

**Fig. 1.**
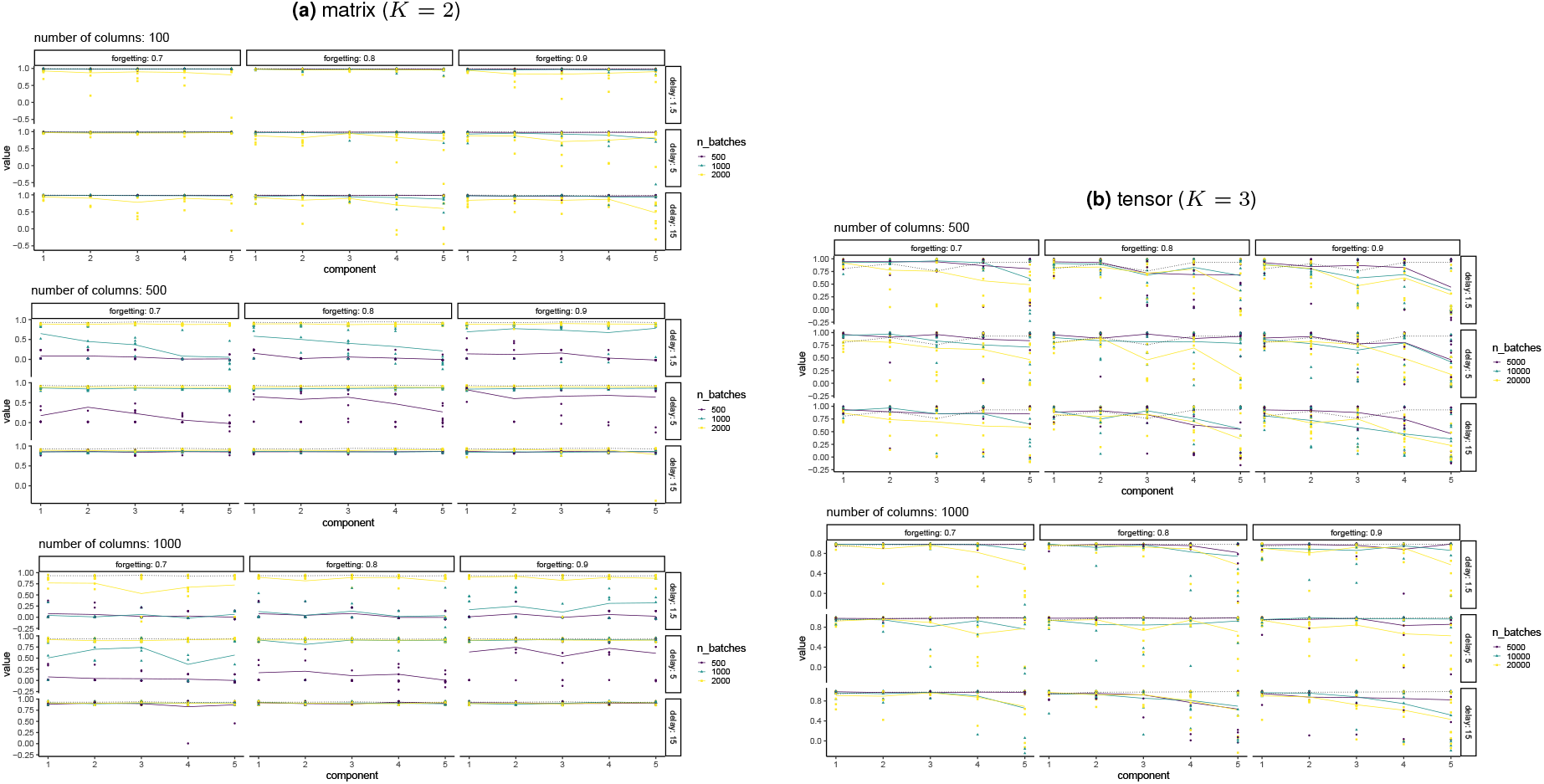
The correlation coefficients between true and estimated parameters *V*. The colored solid line indicates sample mean of the estimates. The dots represent each estimate across all trials. The dotted lines stands for results of batched procedure.`n_batches` stands for size of mini-batches.

Overall, the simulation study demonstrates two key methodological advantages. First, by expressing SVB updates solely in terms of nonzero entries and index distributions, the framework achieves scalability without sacrificing probabilistic rigor. Second, the skip-zeros property allows the use of very large mini-batches, which reduces variance and improves estimation while avoiding the memory limitations that plague conventional approaches. Together, these results validate both the statistical soundness and computational efficiency of UNISON, confirming its suitability for large-scale single-cell analysis in the million-cell era.

### Large-scale single-cell analysis of the MOCA dataset

To demonstrate the scalability and practical utility of the skipzeros SVB framework, we applied it to one of the largest publicly available single-cell transcriptomic resources: the Mouse Organogenesis Cell Atlas (MOCA) dataset (11). This dataset profiles more than one million cells across mouse embryonic development, thereby providing an ideal benchmark for assessing both computational performance and biological interpretability. The MOCA count matrix comprises 26,183 genes measured across 1,331,984 cells, with only 2.6% of entries being nonzero. Such extreme sparsity poses a major challenge for conventional NMF implementations, which typically expand zeros in memory and thus become intractable at this scale. In contrast, our skip-zeros SVB framework directly operates on the COO sparse representation, leveraging only nonzero entries for computation. For benchmarking, we compared our method with Liger’s online NMF algorithm (7), which has been widely applied for large-scale single-cell integration. Although Liger is memory-efficient, it samples subsets of the data at each iteration and optimizes mean squared error (MSE), which is equivalent to maximizes a Gaussian likelihood and may be less appropriate for highly discrete single-cell count data. In our analysis, we set the latent dimension to *L* = 30, corresponding to the dimensionality used in the original PCA-based preprocessing of MOCA. For our method, we used a mini-batch size of 10^8^, a delay parameter *τ* = 15, and a forgetting rate *κ* = 0.8. The results (Table 1) illustrate both the strengths and trade-offs of our framework. Memory usage was greater than that of Liger (17.7 GB versus 0.63 GB), as our method explicitly incorporates all nonzero entries. However, this increase is dramatically smaller compared to the dense matrix expansion required by conventional NMF (which would theoretically reach hundreds of gigabytes for this dataset), demonstrating the effectiveness of the skip-zeros strategy. Computation time remained practical, requiring approximately 9.8 hour for 100 epochs over the full dataset.

**Table 1.**
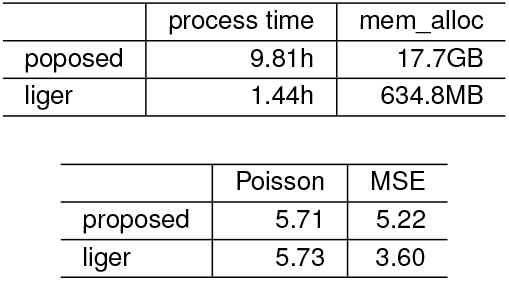
Benchmark UNISON vs. rliger. mem_alloc stand for total amount of memory allocated.

|  | process time | mem_alloc |
| --- | --- | --- |
| poposed | 9.81h | 17.7GB |
| liger | 1.44h | 634.8MB |

|  | Poisson | MSE |
| --- | --- | --- |
| proposed | 5.71 | 5.22 |
| liger | 5.73 | 3.60 |

More importantly, the latent factors estimated by our method yielded superior biological interpretability. When cell embeddings inferred from the factor matrix *V* ^(2)^ were projected into UMAP space (Figure 2), cell populations corresponding to distinct developmental lineages were more clearly separated compared to those obtained with Liger. This suggests that explicitly modeling count distributions with a Poisson likelihood produces latent factors that more faithfully capture biologically meaningful variation.

**Fig. 2.**
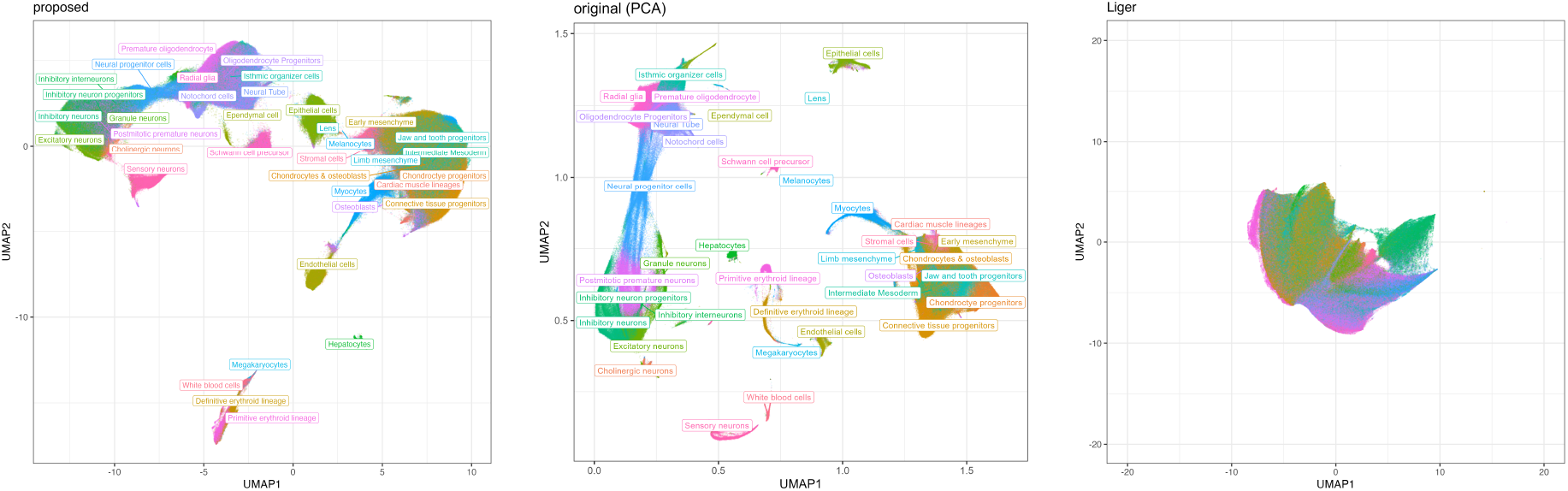
Summary of MOCA data using NMF. Left: UMAP representation of cell features *V* ^(2)^. Center: Original UMAP representation in Cao et al’s study. Right: UMAP representation of cell features by Liger which corresponds to liger in Table 1.

The nonnegative constraints inherent in NMF further encouraged sparsity in the latent representation, resulting in components enriched for lineage-specific marker genes. These factors aligned well with known developmental trajectories in embryogenesis. By contrast, Liger’s Gaussian model tended to produce diffuse components that were less easily interpretable. Thus, the combination of probabilistic modeling and skip-zeros optimization not only enabled scalability to more than one million cells but also enhanced the ability to extract biologically relevant latent structure. In summary, the analysis of the MOCA dataset demonstrates the core utility of UNISON, which enables decomposition of more than one million cells and highlights its applicability to the million-cell scale of modern single-cell transcriptomics. The framework achieves a balance between computational feasibility and interpretability, allowing large-scale single-cell datasets to be decomposed in a statistically principled manner. By avoiding explicit expansion of zero entries, the framework provides a practical solution for analyses that were previously computationally prohibitive, and it recovers latent factors that capture both global structure and lineage-specific signatures in developmental processes.

To examine each component of the latent variable *V*, we calculated the frequency-exclusivity (frex; 8) score. Here, frex is defined as the harmonic mean of the two probabilities *p*(*r*|*l*) and *p*(*l*|*r*), where *r* denotes a cell type or pathway and *l* denotes the component index. The results are shown in Figure 3. The frequency of latent classes (component index) varies across developmental stages (Figure 3a). In the early stages (E9.5–E10.5), components 11 and 17 are relatively abundant, whereas components 26, 24, and 15 become more dominant in the later stages (E12.5–E13.5), indicating stage-specific activity of particular modules. “Excitatory neurons” and “Inhibitory neurons” are prominent in component 17, indicating that it represents neuronal cell classes (Figure 3b). This component also features the “Long-term potentiation” pathway (Figure 3c), suggesting that it corresponds to a module associated with neuronal maturation. Component 15 is characterized by “Definitive erythroid lineage cells,” and “Stromal cells,”(Figure 3b). These populations likely reflect hematopoietic lineages and their supporting mesenchymal environment during the establishment of the hematopoietic niche. Furthermore, component 25 is characterized by “Early mesenchymal cells”, “Jaw and tooth progenitors”, and “Chondrocyte progenitors” (Figure 3b). These populations represent craniofacial mesenchymal lineages and are likely involved in skeletal and cartilage development during early morphogenesis. Overall, these latent classes are consistent with known developmental stages, cell types, and pathways, suggesting that NMF successfully extracts biologically meaningful features.

**Fig. 3.**
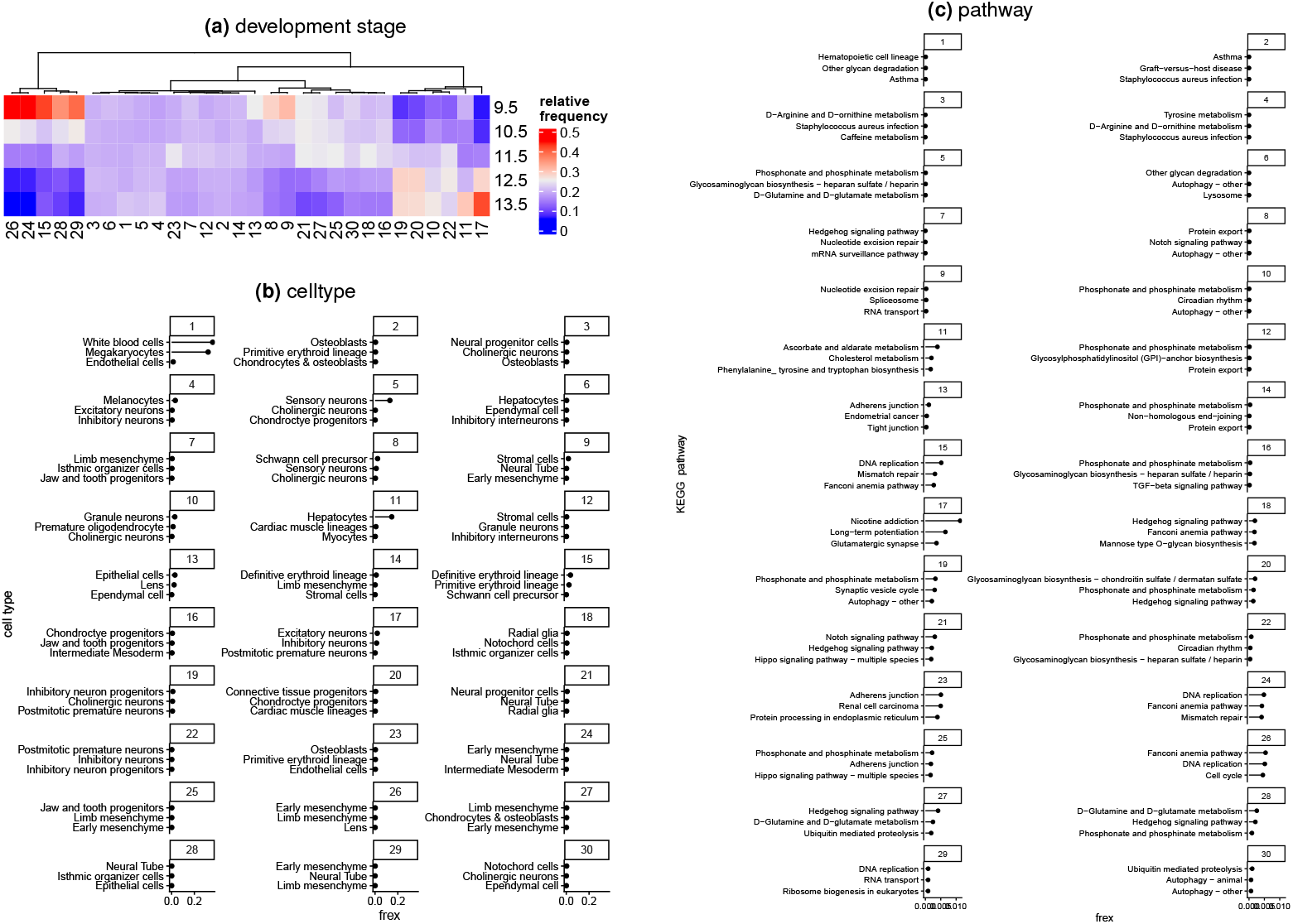
Characterization of latent clusters across developmental stage, cell type, and functional pathways. For cell type and pathway, the three with the largest frex values were displayed. (a) Relative frequencies of each latent cluster along the developmental timeline. Here, y-axis stands for the frequency which normalized by each development stage. (b) Cell types characterizing each cluster. (c) KEGG pathway characterizing each cluster.

### Cross-species single-cell analysis

To further illustrate the flexibility of the skip-zeros SVB framework, we applied it to the cross-species single-cell transcriptomic dataset of across the multiple life stages for mice, zebrafish and *drosophila* (12). This source profiles over two million cells.

To enable joint analysis, we constructed a design matrix *Z* encoding categorical variables including species identity, sample, cell, and gene information. Homologous genes (as shown in Table 2) were mapped across species, thereby embedding all cells into a shared latent space. The UNMF formulation is particularly well suited to this setting, as it separates the format of sparse data from contextual variables. By combining this with skip-zeros SVB updates, we ensured that the inference procedure scaled efficiently to the size of the dataset without the need to enumerate zero entries. In this analysis, we set the latent dimension to *L* = 30, a mini-batch size of 10^7^, a delay parameter *τ* = 15, and a forgetting rate *κ* = 0.8.

**Table 2.** Example of the corresponding the genes between different species. Top table convert to bottom table. Here, the tables stand for the homologous gene. However, in consideration of other possible applications, it can be understood as an auxiliary information to the row and/or column of matrix.

| gene | mouse gene |  |  |  |
| --- | --- | --- | --- | --- |
| zyg11 | Zyg11a, Zyg11b |  |  |  |
| zyx | Zyx |  |  |  |
| ⋮ | ⋮ |  |  |  |
| ⋮ | ⋮ |  |  |  |
|  | ↓ |  |  |  |
|  | Zyg11a | Zyg11b | Zyx | ⋯ |
| zyg11 | 1 | 1 | 0 |  |
| zyx | 0 | 0 | 1 |  |
| ⋮ | ⋮ | ⋮ | ⋮ | ⋯ |

Figure 4 presents selected estimation results for *V*. The latent variables for each cell do not differ clearly between species (Figure 4a), whereas the latent variables for each sample tend to differ across species (Figure 4b). This pattern suggests that species-specific differences are largely absorbed by the sample-level factors, and thus may not appear explicitly in the cell-level factors.

**Fig. 4.**
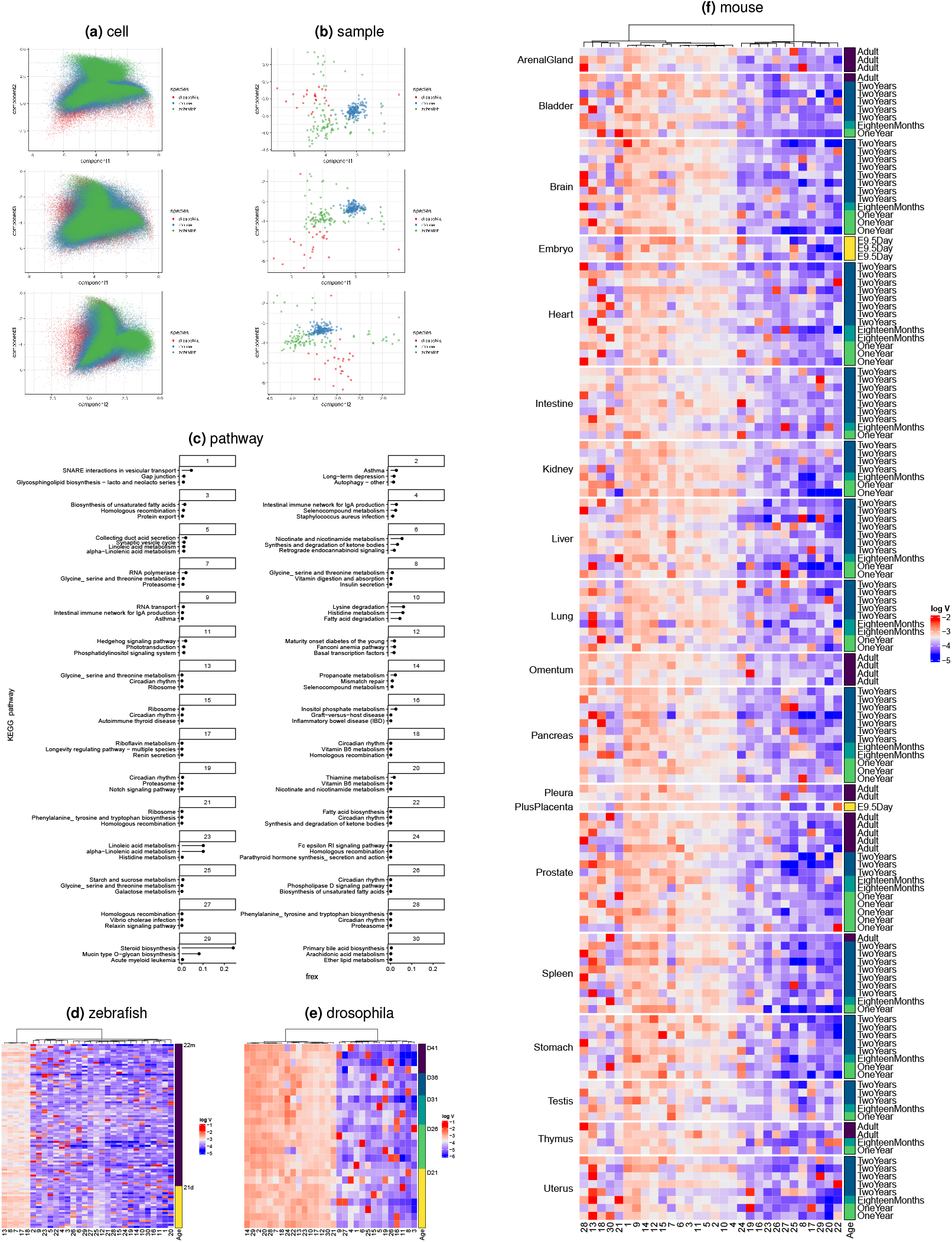
Characterization of latent components across developmental stage, cell type, and functional pathways. (a) scatter plots for cell factors (b)scatter plots for sample factors (c) Homologous genes that were set as common factors for all species were displayed. (d-f) Heatmap of sample factors (d) zebrafish (e) drosophila (f) mouse.

Next, we show the mouse homologous genes that we designated as common factors (Figure 4c-f). In a simplified summary, Wang et al. reported increases in immune-related pathways and decreases in metabolic pathways. Component 20 is associated with “Thiamine metabolism,” “Vitamin B6 metabolism,” and “Nicotinic acid and nicotinamide metabolism”—that is, vitamins B1, B3, and B6—and is interpreted as a module involved in adenosine triphosphate (ATP) production. The components on the right side of Figure 4d, such as components 8 and 20, tend to increase with age. This observation is consistent with Wang et al., although some organs exhibit the opposite trend (e.g., the kidney in Figure 4d). The ability to summarize such complex, large-scale data and provide interpretable results for analysts is one advantage of the proposed method.

Finally, for the zebrafish and Drosophila genes designated as species-specific factors, the KEGG pathways most closely related to each component are shown in Figure 5. Components 17 and 18 appear in pathways with high frex values in both zebrafish and Drosophila, although their functional interpretations differ. In zebrafish, component 17 is associated with “Autoimmune thyroid disease,” “Hedgehog signaling pathway,” and “Intestinal immune network for IgA production,” suggesting a role in immune function. In contrast, in Drosophila, component 18 is inferred to play an immune role because it is involved in “African trypanosomiasis,” “Retinol metabolism,” and “D-glutamine and D-glutamate metabolism.” These represent species-specific gene expression patterns that cannot be explained by the common components.

**Fig. 5.**
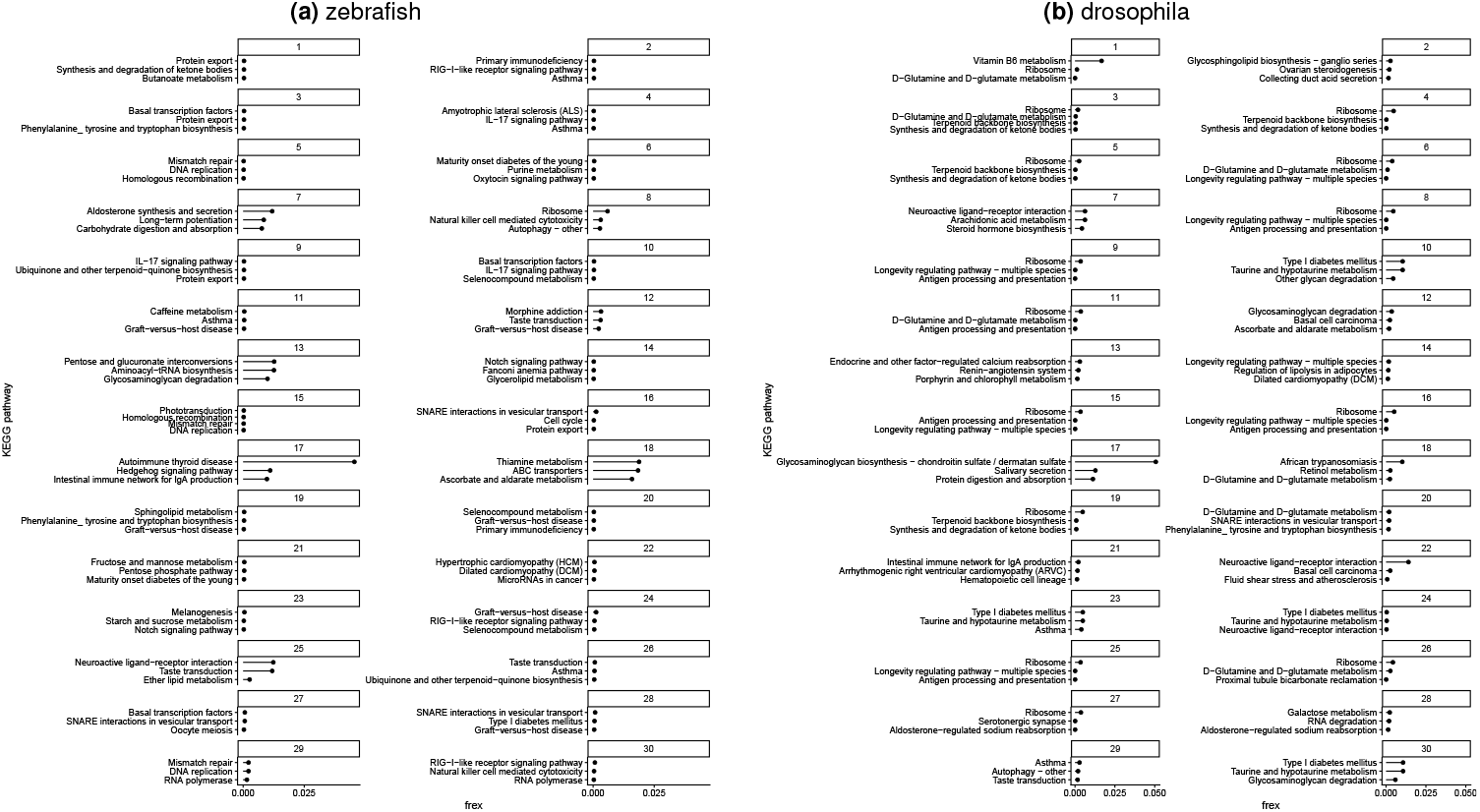
Characterization of latent components via functional pathways enriched in species-specific gene. (a) and (b) are specific to zebrafish and Drosophila, respectively.

Taken together, this cross-species analysis highlights two major advantages of UNISON, which extends the skip-zeros SVB principle to heterogeneous multi-species data integration. First, by leveraging the count-based Poisson likelihood while skipping zeros in computation, the method scales efficiently to large and heterogeneous datasets. Second, by incorporating contextual information through the design matrix, the model disentangles species-specific variation from conserved programs, yielding interpretable latent factors that align with known biological functions. In this way, the frame-work supports integrative analysis across multiple species, thereby broadening the scope of single-cell transcriptomic studies.

## Discussion

In this study, we introduced UNISON, a scalable framework for nonnegative matrix factorization and its extensions, based on skip-zeros variational inference. This innovation directly addresses the computational challenges of the million-cell era of single-cell transcriptomics. By reformulating variational updates in terms of sufficient statistics, we demonstrated that inference can be performed without explicit access to zero entries in sparse matrices. This innovation provides both a theoretical guarantee of statistical correctness and a practical route to analyzing single-cell datasets at unprecedented scale.

Compared to existing approaches, our framework offers several advantages. Online NMF methods (13, 14) achieve scalability through subsampling, but at the cost of discarding substantial fractions of the data and optimizing objectives that may be less well matched to discrete count distributions. Our skip-zeros SVB algorithm, by contrast, incorporates all observed nonzero counts into the estimation procedure, while accounting for zeros implicitly through geometric sampling. This preserves the statistical fidelity of the underlying generative model while ensuring computational efficiency. Furthermore, unlike nonlinear embedding methods such as UMAP or t-SNE, which prioritize visualization but yield non-deterministic representations, matrix factorization provides reproducible, linearly structured latent factors that can be directly incorporated into downstream probabilistic analyses. The results on both simulated and real data confirm the practical advantages of our approach. In simulations, we demonstrated robustness across a wide range of learning-rate schedules and mini-batch sizes, and highlighted clear guidelines for parameter selection in practice. Analysis of the MOCA dataset showed that the method scales to over one million cells, yielding latent factors that not only capture global developmental trajectories but also reveal lineage-specific signatures with greater interpretability than competing approaches. The cross-species application further highlighted the flexibility of the framework: by incorporating background information such as species identity through the UNMF design matrix, the model was able to disentangle conserved from species-specific transcriptional programs, and to recover biologically meaningful gene-gene and gene-phenotype associations relevant to glaucoma.

While the results are promising, several limitations merit discussion. First, although the skip-zeros SVB framework substantially reduces memory usage, large-scale applications still require nontrivial computational resources, particularly when applied to datasets containing hundreds of millions of cells. Second, the current implementation relies on the Poisson likelihood, which may not fully capture overdispersion or zero-inflation frequently observed in single-cell RNA-seq data. Extensions to negative binomial or zero-inflated models (15, 16) could further improve robustness in these settings. Third, while the geometric sampler provides an unbiased approximation to the contribution of zeros, alternative strategies for variance reduction or control variates may accelerate convergence.

Despite these limitations, our work provides a general and extensible foundation for large-scale single-cell analysis. By combining statistical rigor with computational efficiency, the skip-zeros SVB framework bridges a critical gap between the growing scale of single-cell experiments and the analytical methods available to interpret them. Beyond single-cell transcriptomics, the principles underlying our approach—efficient exploitation of sparsity, probabilistic factorization, and integration of contextual variables—are broadly applicable to other high-dimensional biological datasets, including epigenomic profiles, proteomics, and multi-omics integration.

In conclusion, we believe that UNISON, powered by skipzeros variational inference, represents a key step toward making truly large-scale and integrative single-cell studies feasible in the million-cell era. As experimental technologies continue to increase the scale and diversity of data generation, methods that exploit sparsity while retaining interpretability will be essential. We expect that UNISON will serve as a foundational methodology for the analysis of high-dimensional, sparse biological data in the years to come.

## Materials and methods

### Data structure for sparse matrices

Single-cell transcriptomic data are typically represented as sparse count matrices, in which the overwhelming majority of entries are zero. To exploit this sparsity, we adopt the COO format, which records only nonzero elements as triplets:

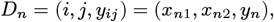

corresponding to the (*i, j*)-th element of the matrix *y*_*ij*_. The complete dataset is then expressed as 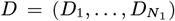, where *N*_1_ denotes the number of nonzero elements. Table 3(top) illustrates this representation. On the left, the matrix is stored in its dense form, with explicit zeros. On the right, the same matrix is expressed in COO format, where only the nonzero elements and their indices are retained. This compact representation drastically reduces memory requirements and eliminates the need to manipulate zeros directly.

**Table 3.**
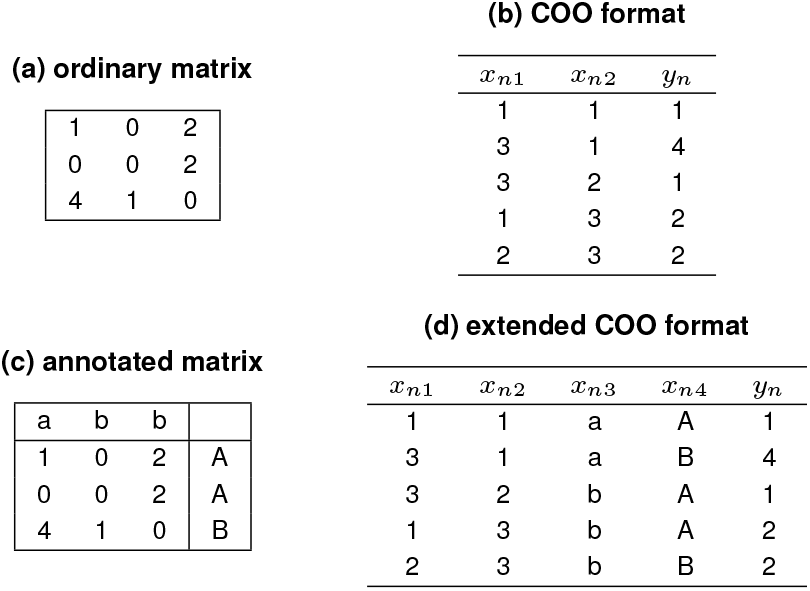
Sparse matrix. The left and right tables have exactly same information, respectively. The form of tables (b) and (d), are used in this study.

This COO format is widely supported in computational biology. It is equivalent to the `TsparseMatrix` class of the R Matrix package (17), the Matrix Market format, and the sparse matrices output by pipelines such as Cell Ranger(18). The COO representation naturally generalizes to higher dimensions. Specifically, each observation can be expressed as

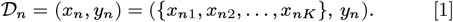

where *x*_*n*_ encodes categorical indices across *K* dimensions (e.g., gene, cell, species, batch, condition), and *y*_*n*_ is the observed count. This formulation allows sparse arrays and annotated matrices to be handled in a unified manner. Table 3 (bottom) provides an example of this extended representation. On the left, the dense matrix includes an additional categorical annotation (A or B). On the right, the COO format encodes this information explicitly as part of the tuple ({*x*_*n*1_, *x*_*n*2_, *x*_*n*3_, *x*_*n*4_}, *y*_*n*_).

This extension demonstrates that the COO representation can incorporate categorical or contextual variables (such as species, sample, or batch) directly into the index set, while still storing only nonzero entries. As such, it provides a unified framework for representing large-scale sparse matrices and tensors, which is crucial for efficient estimation in subsequent analyses.

### Evaluation of the likelihood with sparse matrices

To perform Bayesian or maximum likelihood estimation of unknown parameters *θ*, evaluation of the likelihood function is required. For models in the exponential family, the likelihood depends only on sufficient statistics. Formally, the exponential family is defined as:

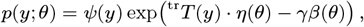

where the tr placed at the upper left of a matrix or vector denotes the transpose, *T* (*y*) is the sufficient statistic, *η*(*θ*) is the natural parameter, and *β*(*θ*) is the log-normalizer (or log-partition function). The scalar *γ* is a known constant (typically the sample size) that does not depend on *y*. The exponential family admits a conjugate prior with hyperparameters *ξ* = (*ξ*_1_, *ξ*_0_):

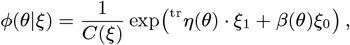

where *C*(*ξ*) is the normalizing constant. Given this conjugate prior, the posterior distribution is:

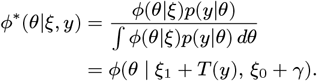

A critical observation is that *β*(*θ*) does not depend on the observed values *y*, while the sufficient statistic *T* (*y*) vanishes for *y* = 0 whenever *T* (0) = ^tr^(0, …, 0). This means that Bayesian updates can be performed without accessing the zero entries of a sparse matrix. In practice, mixtures of exponential family distributions do not themselves belong to the exponential family. However, variational Bayes (VB; 19, 20) with a mean-field approximation can still be applied due to the conditional conjugacy of exponential family priors. Let *y*_*n*_; (*n* = 1, …, *N*) denote exponential-family random variables that are conditionally independent given indices *x*_*n*_ = ^tr^(*x*_*n*1_, …, *x*_*nK*_). The log-likelihood of parameter *θ*_*k*_, corresponding to the *k*-th axis, is:

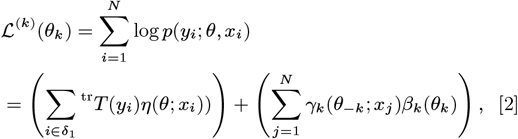

where *θ*_*−k*_ denotes all parameters except *θ*_*k*_, and *δ*_1_ is the set of indices corresponding to nonzero elements. In the case of an ordinary matrix with *K*^(1)^ rows and *K*^(2)^ columns, *N* = *K*^(1)^ · *K*^(2)^. Importantly, the first term depends only on the nonzero entries, whereas the second term depends only on indices, not on the observed values *y*. Thus, if the index ranges are known, the log-likelihood can be evaluated without accessing zero elements. To implement stochastic gradient or variational methods, we typically resample from the empirical distribution. Let *s* denote a random index, and *L*_*s*_(*θ*) the log-likelihood contribution from the sampled element *D*_*s*_. The expectation of the log-likelihood is then:

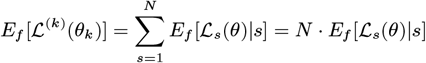

where let *E*_*f*_ [·] be the expectation by the empirical distribution. If 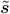 is sampled uniformly from 1, …, *N*_1_, i.e. only from nonzero entries, we can approximate:

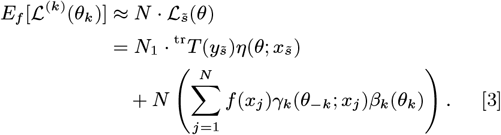

When *y*_*i*_ = 0, the first term vanishes, so zero entries are never required. The second term depends only on the index distribution *x*_*j*_, which is known a priori. As a further extension, consider a stochastic mini-batch *S*_*m*_ of size 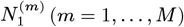. The expectation of the log-likelihood is then:

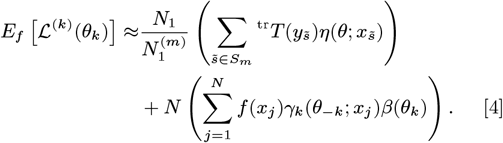

Equation [4] demonstrates that both stochastic gradient ascent (SGA) and SVB can be realized without ever accessing zero elements. This constitutes the central result of this subsection and establishes the theoretical foundation for UNISON developed below.

### Generic SVB algorithm for sparse matrix factorization

An important special case of the general framework is ordinary matrix factorization, where *K* = 2. Using the notation introduced in Equation [1], the observed count matrix *Y* can be approximated as the product of two low-rank factor matrices.

Under the mean-field assumption, the variational posterior for *θ* = (*U, V* ^(1)^, *V* ^(2)^) factorizes as:

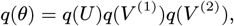

where *U* denotes auxiliary latent variables, and *V* ^(1)^ and *V* ^(2)^ correspond to the row- and column-specific factors, respectively. The expectation of the log-likelihood with respect to *q*(*θ*\*V* ^(*i*)^) is:

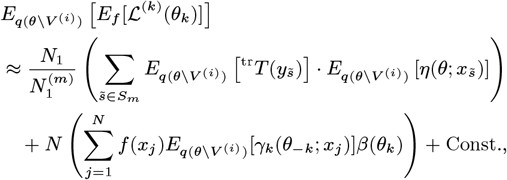

Here, Const. represents terms that do not depend on *V* ^(*i*)^. Since the empirical distribution of indices is uniform in many practical cases (e.g., each row has the same number of columns), explicit calculation of *f* (*x*) is unnecessary. In contrast, if the matrix has random missing values, we can handle it using *w*_*i*_ = *N* · *f* (*x*_*i*_). The variational posterior update for *V* ^(*i*)^ at iteration *t* is:

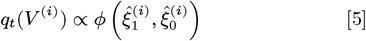

with variational parameters:

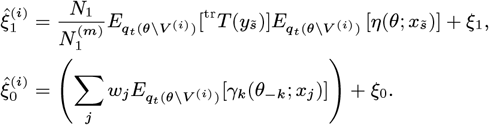

#### Algorithm 1 SVB procedure with sparse matrix. Let 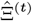 be the all of the variational parameters at step *t*.

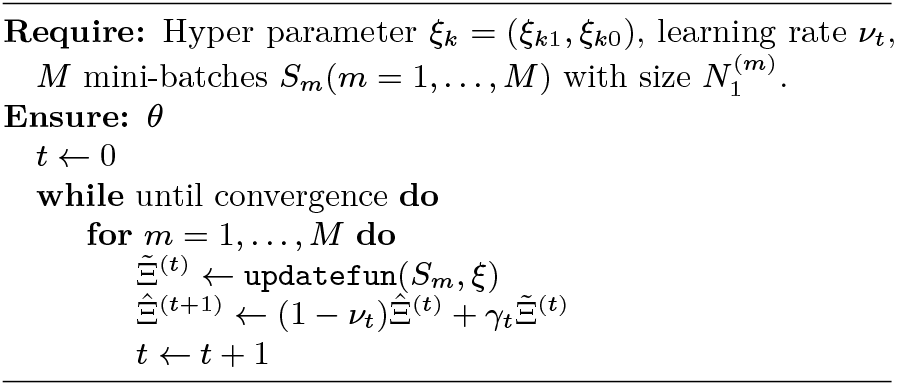

### Learning rate schedule

The learning rate *v*_*t*_ follows an exponential decay:

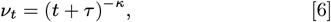

where smaller *κ* yields faster but less stable updates, and larger *τ* slows the decay, ensuring stable convergence in large-scale datasets. This generic SVB algorithm provides the foundation for efficient Bayesian inference in NMF with sparse matrices, and serves as a core component of UNISON. The specific form of the updates for Poisson likelihood is presented in the following subsection.

### Generic SVB algorithm for regression models with categorical predictors

The skip-zeros SVB framework can be extended beyond standard matrix factorization to more general regression problems of the form *p*(*y*|*x*_*n*_), where *x*_*n*_ encodes categorical predictors. This extension provides the theoretical foundation for UNMF, which allows simultaneous analysis of multiway and context-dependent data structures.

#### Reformulation of the log-likelihood

Using the notation introduced in Equation [1], matrix factorization is a special case of regression, where *x*_*n*_ = (*x*_*n*1_, …, *x*_*nK*_) indexes the predictors, and *y*_*n*_ follows an exponential family distribution. To avoid explicit handling of zeros, we assume independence among the predictor variables *X*. Under this assumption, Equation [3] can be decomposed as:

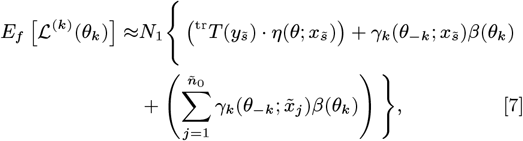

where *ñ*_0_ is the random number of zero elements (“failures”) observed before encountering a nonzero element (“success”), which follows a geometric distribution. The indices 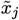 are drawn accordingly. Thus, the contribution of zero elements can be approximated stochastically via random sampling rather than explicit enumeration.

#### Mini-batch formulation

In practice, we employ stochastic mini-batches of nonzero entries. For a mini-batch *S*_*m*_ of size 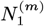, the expectation of the log-likelihood becomes:

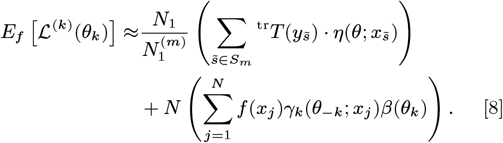

The first two terms depend only on observed nonzero entries, while the third term represents the expected contribution of zero entries, approximated using geometric sampling. This formulation ensures that SVB updates can be performed using only mini-batches of nonzero entries, while faithfully incorporating the statistical influence of zeros.

#### Implications for UNMF

Equation [4] shows that the skip-zeros SVB framework generalizes naturally to regression models with categorical predictors. This insight enables efficient estimation in UNMF, where categorical variables (e.g., species, condition, cell type) are encoded in a design matrix and incorporated directly into the factorization model. The estimation problem is thereby reduced to computing expectations of the sufficient statistics *T* (*y*), the natural parameter *η*(*θ*), and the log-partition function *β*(*θ*),all of which can be evaluated without explicit enumeration of zeros. This reformulation establishes the theoretical basis for UNISON, which extends skip-zeros SVB to unified NMF and supports integrative analyses across heterogeneous conditions, as described in the following section.

### Matrix factorization and its extension with Poisson loss

We now consider an important special case of the generic frame-work: Poisson nonnegative matrix factorization (Poisson NMF). This formulation is particularly suitable for single-cell RNA-seq data, where observations are nonnegative counts and sparsity is extreme.

#### Poisson NMF model

Let *X* be a nonnegative integer matrix of size *D*_1_ × *D*_2_. The objective is to approximate *X* by the product of two nonnegative factor matrices:

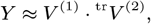

where 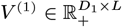 and 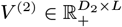, and *L* is the latent dimension (rank of the decomposition). In the context of single-cell analysis, *V* ^(1)^ represents latent factors for gene features, while *V* ^(2)^ represents latent factors for cell features. Because scRNA-seq data are counts, the Poisson likelihood is a natural choice. Indeed, minimizing the generalized Kullback-Leibler (KL) divergence commonly used in NMF (2) is equivalent to maximizing the Poisson likelihood. Following widely used formulations(3), the generative process is expressed as:

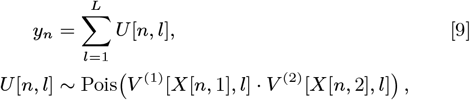

with Gamma priors imposed on the factor matrices:

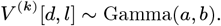

#### SVB updates for Poisson NMF

Using the sufficient-statistic formulation from Equation [3], the stochastic variational Bayes procedure for Poisson NMF with skip-zeros can be summarized as Algorithm 2.

The procedure can be extended to higher-order factorizations with *K* ≥ 3. In practice, however, general multiway relationships are often semi-paired or heterogeneous (e.g., across species, cell types, or conditions). In such cases, UNMF offers a more flexible formulation. By introducing a design matrix *Z*—for example, using one-hot encoding of categorical variables—UNMF can represent multi-dimensional relationships in a consistent framework. Indeed, the UNMF model (8) encompasses Equation [9] as a special case. The SVB algorithm implemented in UNISON, which builds on this formulation to generalize Poisson NMF to heterogeneous contexts, is described in the following section.

### UNISON: Unified Sparse-Optimized Nonnegative factorization

We define UNISON, which extends the skip-zeros SVB frame-work to UNMF across multiple categorical predictors and higher-order relationships.

#### UNMF model formulation

Let *Z* denote a design matrix constructed via one-hot encoding of categorical variables such as species, condition, or sample. The UNMF model is then defined as:

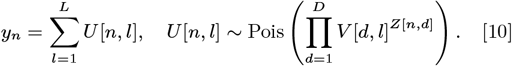

When *Z* is transformed from two categorical variables with one-hot encoding, formulation reduces to the standard Poisson NMF model (Equation [9]).

#### SVB updates for UNMF with skip-zeros

The SVB update procedure for UNMF is summarized in Algorithm 3. As in Poisson NMF, only nonzero entries need to be explicitly accessed. The contribution of zeros is incorporated using a geometric sampler, which approximates their expected effect without enumerating all zero entries.

#### Geometric sampler

The use of geometric sampling ensures that the expected contribution of zero entries can be incorporated without explicitly iterating over them. This dramatically improves scalability, as enumerating all zeros would be computationally infeasible in large-scale single-cell datasets. By combining the COO sparse representation with variational updates, the UNMF with skip-zeros provides a unified and computationally efficient framework for integrating heterogeneous datasets. For example, homologous genes across species can be encoded as shared predictors in *Z*, allowing cells from different species to be embedded into a common latent space. In this way, UNISON supports cross-species or multi-condition single-cell analysis while retaining interpretability and computational feasibility.

## ACKNOWLEDGMENTS

The Grant-in-Aid for Transformative Research Areas (platforms for Advanced Technologies and Research Resources) (grant no. 22H04925) and Grant-in-Aid for Transformative Research Areas (A) (grant no. 23H04938) were provided by the Japan Society for the Promotion of Science (JSPS). Additional support for T.S. was received from the Japan Agency for Medical Research and Development (AMED) through the Core Research for Evolutional Science and Technology (grant no. JP25gm2010002), the Project Promoting Support for Drug Discovery (grant no. JP25nk0101112), Brain/MINDS Health and Diseases (grant no. JP25wm0625519), the Interdisciplinary Cutting-edge Research (grant no. JP25wm0325068), the Moonshot R&D Program (grant no. JP25zf0127012), and the Advanced Genome Research and Bioinformatics Study to Facilitate Medical Innovation (GRIFIN) (grant no. JP25tm0424226). Further funding for T.S. was provided by the Japan Science and Technology Agency (JST) under the Moon-shot R&D program (grant no. JPMJMS2025). Y.K. was supported by the Project for P-PROMOTE (grant no. 24ama221609h0001) from AMED, the National Cancer Center Research and Development Fund (grant no. 2024-A-6), and a JSPS Grant-in-Aid for Early-Career Scientists (grant no. 23K16991). Further support came from the Medical Research Center Initiative for High Depth Omics and Multilayered Stress Diseases at the Institute of Science Tokyo. Supercomputing resources were provided by the Shirokane supercomputer at the Human Genome Center, the University of Tokyo, and the TSUBAME3.0 supercomputer at the Institute of Science Tokyo.

## Algorithm 2 SVB procedure for Poisson NMF with skipping zeros

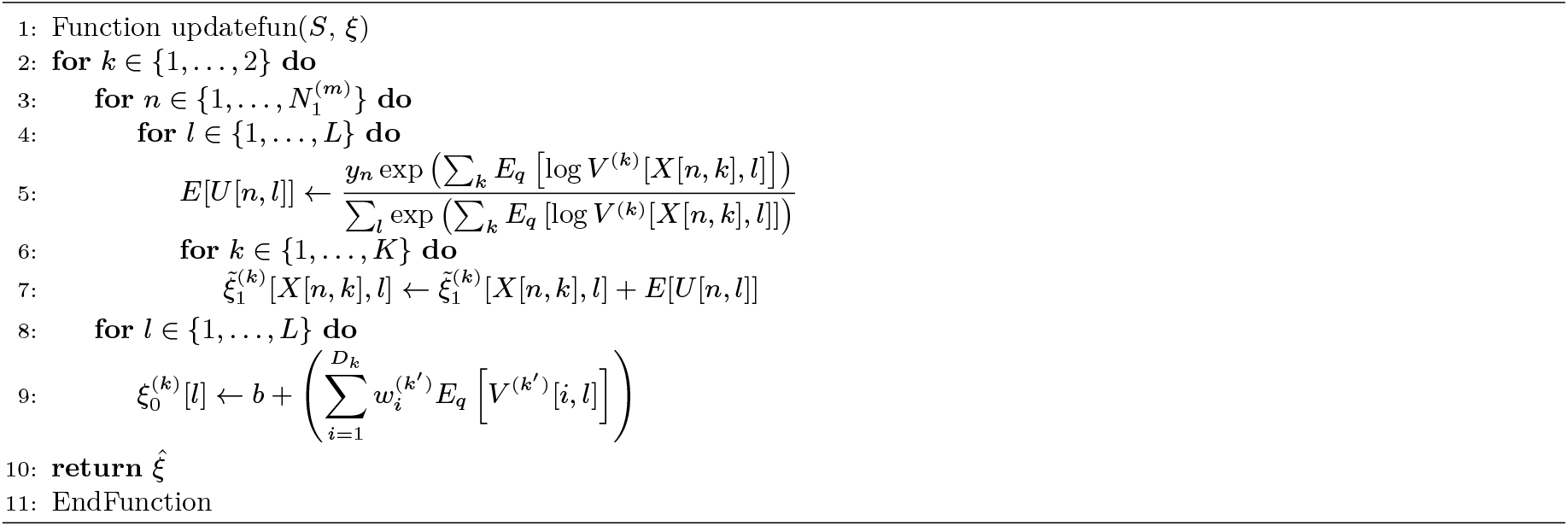

## Algorithm 3 SVB procedure for UNMF with skipping zeros. geometric_sampler is sampling procedure for variational parameter *ξ*_0_. It’s shown in Algorithm 4.

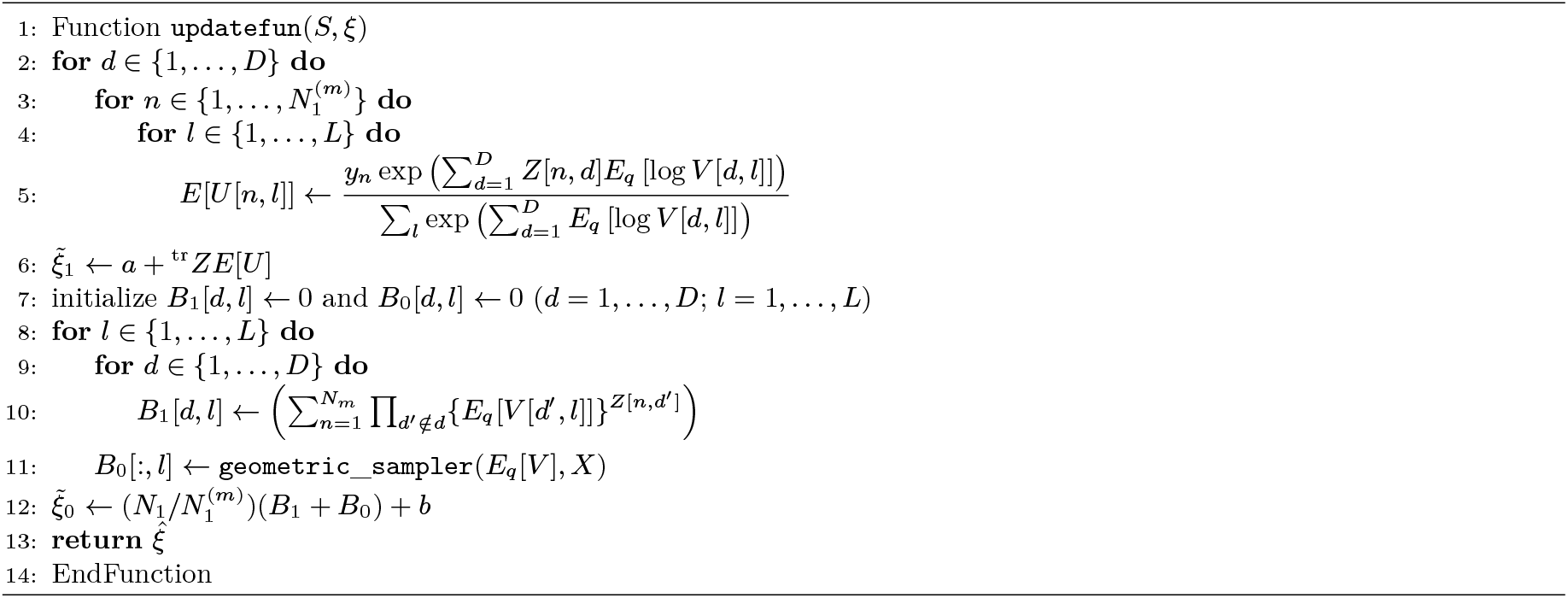

## Algorithm 4 SVB procedure for UNMF with skipping zeros. geometric_sampler is sampling procedure for variational parameter *ξ*_0_. Its corresponds to in the third term of Equation [8].

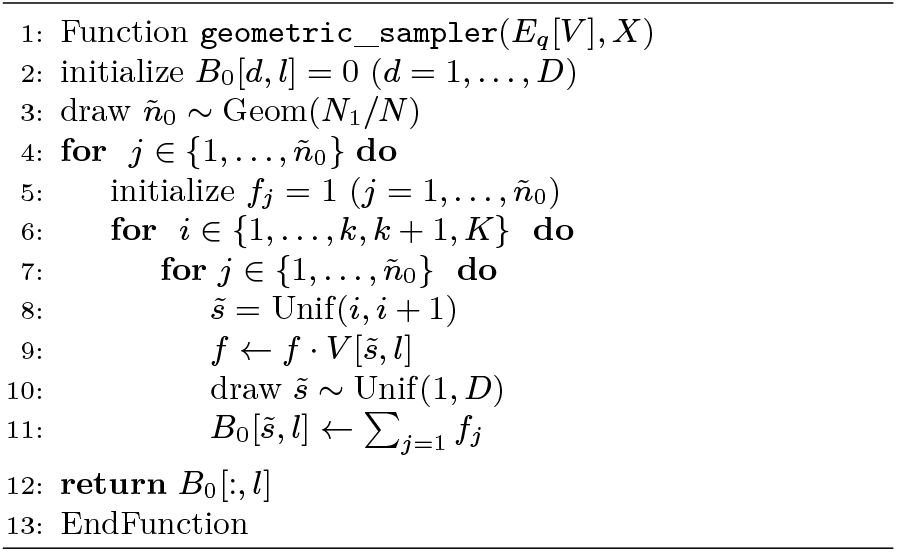

## References

1. S Lobato-Moreno, et al., Single-cell ultra-high-throughput multiplexed chromatin and rna profiling reveals gene regulatory dynamics. Nat. Methods 22, 1213–1225 (2025).

2. D Lee, HS Seung, Algorithms for non-negative matrix factorization. Adv. neural information processing systems 13 (2000).

3. AT Cemgil, Bayesian inference for nonnegative matrix factorization models. Comput. Intell. Neurosci. p. 785152 (2009).

4. A Diaz-Papkovich, L Anderson-Trocmé, C Ben-Eghan, S Gravel, Umap reveals cryptic population structure and phenotype heterogeneity in large genomic cohorts. PLoS genetics 15, e1008432 (2019).

5. L Van der Maaten, G Hinton, Visualizing data using t-sne. J. machine learning research 9 (2008).

6. T Chari, L Pachter, The specious art of single-cell genomics. PLOS Comput. Biol. 19, e1011288 (2023).

7. R Gaujoux, C Seoighe, A flexible r package for nonnegative matrix factorization. BMC bioinformatics 11, 1–9 (2010).

8. K Abe, T Shimamura, UNMF: a unified nonnegative matrix factorization for multi-dimensional omics data. Briefings Bioinforma. 24, bbad253 (2023).

9. MD Hoffman, DM Blei, C Wang, J Paisley, Stochastic variational inference. J. Mach. Learn. Res. (2013).

10. E Angelino, MJ Johnson, RP Adams, Patterns of scalable Bayesian inference. Foundations Trends Mach. Learn. 9, 119–247 (2016).

11. J Cao, M Spielmann, X Qiu,, et al., The single-cell transcriptional landscape of mammalian organogenesis. Nature 566, 496–502 (2019).

12. R Wang, et al., Construction of a cross-species cell landscape at single-cell level. Nucleic acids research 51, 501–516 (2023).

13. B Cao, et al., Detect and track latent factors with online nonnegative matrix factorization. in IJCAI. Vol. 7, pp. 2689–2694 (2007).

14. R Zhao, VY Tan, Online nonnegative matrix factorization with outliers. IEEE Transactions on Signal Process. 65, 555–570 (2016).

15. H Abe, H Yadohisa, A non-negative matrix factorization model based on the zero-inflated tweedie distribution. Comput. Stat. 32, 475–499 (2017).

16. O Gouvert, T Oberlin, C Févotte, Negative binomial matrix factorization. IEEE Signal Process. Lett. 27, 815–819 (2020).

17. D Bates, M Maechler, M Jagan, Matrix: Sparse and dense matrix classes and methods (2024).

18. GXY Zheng,, et al., Massively parallel digital transcriptional profiling of single cells. Nat. Commun. 8, 1–12 (2017).

19. MI Jordan, Z Ghahramani, TS Jaakkola, LK Saul, An introduction to variational methods for graphical models. Mach. Learn. 37, 183–233 (1999).

20. DM Blei, A Kucukelbir, JD McAuliffe, Variational Inference: A Review for Statisticians. J. Am. Stat. Assoc. 112, 859–877 (2017).

